# PeTriBERT : Augmenting BERT with tridimensional encoding for inverse protein folding and design

**DOI:** 10.1101/2022.08.10.503344

**Authors:** Baldwin Dumortier, Antoine Liutkus, Clément Carré, Gabriel Krouk

**Affiliations:** Zenith team LIRMM, CNRS UMR 5506 Univ. Montpellier, France; BionomeeX Montpellier, France; Systems, IPSiM Univ. Montpellier, CNRS, INRAE Montpellier, France

**Keywords:** inverse folding, AlphaFold, proteins, protein generation, transformers, NLP

## Abstract

Protein is biology workhorse. Since the recent break-through of novel folding methods, the amount of available structural data is increasing, closing the gap between data-driven sequence-based and structure-based methods. In this work, we focus on the inverse folding problem that consists in predicting an amino-acid primary sequence from protein 3D structure. For this purpose, we introduce a simple Transformer model from Natural Language Processing augmented 3D-structural data. We call the resulting model PeTriBERT: Proteins embedded in tridimensional representation in a BERT model. We train this small 40-million parameters model on more than 350 000 proteins sequences retrieved from the newly available AlphaFoldDB database. Using PetriBert, we are able to *in silico* generate totally new proteins with a GFP-like structure. These 9 of 10 of these GFP structural homologues have no ressemblance when blasted on the whole entry proteome database. This shows that PetriBert indeed capture protein folding rules and become a valuable tool for *de novo* protein design.

## I. Introduction

Proteins are at the heart of cellular life and therefore are the center of numerous medical and chemical applications. They are macro molecules that are built as various length chains composed as combinations of the 20 amino acids (also called residues) found in nature. Proteins fold into 3D conformations depending on their primary sequence and post translational modifications, which define most of their chemical and mechanical properties [1]. Decoding the relationship between the primary sequence of proteins and their structural functions is an important topic of computational biology and has been a long standing problem that was traditionally approached via physical modeling [2]. However, thanks to the recent advent of deep learning and to the growth of both labeled and unlabeled protein databanks, a new range of algorithms arose that operated a paradigm shift in computational biology, leading to unprecedented results [3].

During the last decade, protein structure-related deep learning methods were investigated, both in supervised and unsupervised settings, to predict and understand the folding of proteins and its subsequent tasks. Notably, the ability of deep neural network to capture complex spatial patterns was considered regarding the proteome for which structural data is available to solve various related biological questions: protein design [4]–[6], [5], protein function prediction [7] or model assessment [8]. However, the scope of these methods was limited because of the intrinsic difficulty and cost to experimentally unravel the full structure of the known proteins sequences. This left most of the known proteome unused. To fill this gap, semi-supervised learning have also been investigated, that take benefit of the large amount of available data provided by the abundant use of inexpensive sequencing technologies. Mostly leveraging natural language processing methods [9]–[12], this setting was able to provide task-agnostic representations that can capture important inherent properties to be further finetuned to a wide variety of tasks.

As a matter of fact, it appears that solving the protein folding problem was the missing link between the structure-based and sequence-based paradigms and was hence considered as one key problem to be solved in computational biology. To a large extent, this long-expected question can now be considered solved, since the groundbreaking introduction of AlphaFold 2 [13] and RoseTTAFold [14]. They both use a dual path transformer architecture that combines evolutionary clustered data (embodied as Multiple Segment Alignment) and graph embedding of the residues interactions constrained by geometrical properties to iteratively solve this complex task. AlphaFold in particular exhibited unrivaled performances on this task at the 14th Critical Assessment of protein Structure Prediction (CASP14) challenge, outperforming its competitors by a large margin. AlphaFold is an *adhoc* model that was crafted specifically for this task and its very specific architecture allows it to achieve much better performance than other out-of-the-box architectures.

Since this achievement, Deepmind has publicly released the AlphaFold database [15], which freely offers AlphaFold estimates for the structures of a daily-increasing number of proteins (350,000 at the time we started our experiments). This highly remarkable contribution opens new perspectives in the computational protein field.

In particular, we are interested here in leveraging this very valuable data to investigate a problem that appears as very important in practice in biology, which is *inverse folding*. It consists in the prediction of amino-acid identity of a given unknown residue when the remaining residues and surrounding structure are known. To achieve this, we identify the core difficulty as designing a model that can generate purely synthetic proteins with a given 3D structure.

To achieve inverse folding, we study the use of a slightly tuned version of a light BERT encoder [16] model that uses only 40 million parameters and that we call PeTriBERT for Protein Embedded in a Tridimensional representation in a BERT model. Compared to BERT, our model uses structure data as an additional input which is encoded using learnable fourier features inspired by [17] in order to distillate the newly available data provided by AlphaFold 2.

Our contributions are twofold:

- We show that, despite being a small model, the proposed PeTriBERT inverse folding model is able to capture AlphaFold data well enough so that it can be used for protein design when coupled with Gibbs sampling without additional learning, in a way inspired by [18], [19].
- We benchmark PeTriBERT results by generating proteins sequences having a very close structure to the avGFP [20] and discuss *in silico* evolution of synthetic proteins.

## II. Related work

Recent work in machine learning applied to proteins data aims to solve different tasks such as protein design [4], [5], [21], model assessment [8], [22], protein to protein interaction [12], [23] and predicting protein chemical and mechanical functions [7]. Regardless of the task to solve, different approaches were investigated regarding what input data neural networks model should learn from.

### A. Sequence-based methods

As primary sequences of amino acids are believed to fully encode the information of proteins, sequenced-based machine learning methods arose naturally, relying notably on Natural Language Processing (NLP) frameworks and self-supervision techniques. Self-supervision has recently become a main topic in artificial intelligence research, allowing to benefit from huge amounts of unlabeled data. In NLP, models are trained on proxy tasks such as predicting missing words in a sentence or predicting the next sentence when input some text. They are then finetuned on smaller datasets for specialization on the target task [24].

This particular type of transfer learning was transposed in biology where early works on this topic [25]–[28] were inspired by the well-known word2vec model [24] and where primary sequences of proteins are simply viewed as sentences. As new and more expressive deep neural architectures arose in the general AI community [16], [29]–[33] and with the exponential growth of protein sequence databases, computational biology has recently strongly benefited from this new paradigm.

Recurrent neural networks for proteins modeling were investigated by works such as Heinzinger *et al*. [34] who adapted the ELMo model [35] derived from an LSTM network and showed that NLP is a viable setting for capturing biological properties. Rives *et al*. [9] explored the use of a high capacity transformer network on 250 M of proteins that is the whole known proteome and outperformed models such as LSTM. While learned purely from sequences, they have shown that their model can generalize and actually captures some of the biochemical properties, biological variations, homology et alignment with protein family. Nambiar *et al*. [12] have adapted the RoBERTa transformer [29] and used BPE tokenization to learn task-agnostic representation obtaining comparable results while lowering training time by a huge factor. Brandes *et al*. [36] proposed ProteinBERT using an additional and novel pretraining stage focused on predicting protein functions. Their approach sets a new state-of-the-art performance on a diverse set of benchmarks while using a smaller and faster model. To understand and interpret the attention effect occuring using such architectures, complementary studies were conducted such as the one from Vig *et al*. [37] to specifically interpret the attention maps learned by transformer during the pretraining phase. They have shown that attention captures high-level structural properties of proteins connecting amino acids that are spatially close in three-dimensional structure.

While the transformer model appears to be the most suited architecture, its limitations were exposed by other works such as Rao *et al*. [38] who proposed a benchmark to evaluate NLP-based semi-supervised learning on 5 tasks, using 3 different networks (reccurent, convolutionnal, attention based). They have shown that while self-supervised pretraining improves performance for almost all models on all downstream tasks, performance also varies significantly depending on the chosen architecture across tasks. Other limitations were demonstrated by works such as Rao *et al*. [39], who proposed a transformer architecture that also incorporates evolutionary data encoded as Multiple Sequence Alignement (MSA), which consists in clusters of proteins that share parts of their primary sequences and are believed to testify on the evolutionary history of a protein structure and chemical properties [40]–[42]. They have shown that using MSA allows transformer to outperform single sequence methods and demonstrated the limitations of the latter to accurately encode evolutionary data. Other MSA-based works have also been proposed [43], [44]. The advantage of exploiting MSA features was also illustrated by AlphaFold [13] and RoseTTAFold [14] that achieved unprecedented results using dual-path transformer-based neural networks architecture where one of the paths uses MSA inputs. However, while MSA-based model have shown superiority to single-sequence based models, some other works such as Elnaggar *et al*. [11] showed that MSA features may very well be dropped in some near feature. They compared several state-of-the-art auto-encoders and auto-regressive models and were able to obtain comparable performance to MSA-based model for the first time. Nevertheless, MSA-based methods are still genererally considered to have an edge over single-sequence methods today and the question of capturing the evolutionnary data with sequence-only models remains open.

### B. Structure-based methods

In parallel to sequence-based methods, a more conventional and slightly older research direction has focused on learning representations directly from the available structural data as a way to bypass the complexity of the protein folding problem. Most of the recent methods on that topic either involves convolutional networks [22], [45]–[47] or graph neural networks [4], [5], [7], [8] thanks to their ability to capture tridimensional patterns or instrincic hierarchy between proteins compounds, respectively. Convolutional neural network have notably been extensively used to predict an affinity score beetween compounds based on binding site [48]–[50]), to assess model quality [22] or protein-protein interaction [51]. Graph neural Networks have also been used as an alternative for similar tasks and brought a different approach on model assessment [8], function prediction [7] and binding affinity [52].

More directly related to our present work, another part of structure based methods have focused on inferring spatial predictive distribution of amino-acid given the structure. This is known as the inverse folding problem (IFP) which is often related to protein design as amino acid sequences can then be reversed-engineered from structural constraints [53]. For example, Zhang *et al*. [46] proposed ProDCoNN to predict residue type given the 3D structural environment around the C*α* atom of a residue, which is repeated for each residue of a protein, using 3D meshing around proteins and convolution. Boomsma *et al*. [54] proposed a specific kind of spherical convolution to predict secondary structures of proteins and to solve the inverse folding problem. Anand *et al*. [55] proposed a convolutional network to predict sequences from structure, and performed generation using Conditional Markov Random Field. Chen *et al*. [56] proposed an hybrid method, using both BLSTM and CNN to process 1D structural properties and 2D distance maps, repsectively, and predict binding-site, protein function prediction and residues propensities. Hermosilla *et al*. [45] have used a variant of a Resnet network [57] to model intrinsic/extrinsic distance (distance along amino acid edge and euclidian distance) applied on several tasks, using both primary sequence and protein structure. Strokach *et al*. [4] have used a deep graph neural network to solve the protein design problem as a constraint satisfaction problem applied on the distance matrix of proteins. Jing *et al*. [6] used geometric vector perceptrons, replacing the use of graph or convolutionnal neural networks by a variant of the standard perceptron that have the desired properties of being invariant through rotations and therefore can encode graph features directly without overfitting.

Regarding novel protein generation however, the inverse folding problem is an intermediary step that is discarded in some other works. In particular, encoder-decoder transformer architecture [5], [21], network hallucination [58], [59], generative adversarial networks [60] and variational auto-encoders [61] were also investigated as end-to-end methods on this task.

### C. New hybrid methods

Until recently structured-based models were limited by the low amount of structural data available whereas the sequenced-based problem complexity was such that numerous efforts were necessary to untangle the *language of life* [62]. However, the recent breakthrough in protein folding is progressively closing the gap and structural data is slowly becoming available for a bigger part of the Proteome. It is a strong trend in very recent works to exploit the newly available data. Concurrently to our present work, Zhang *et al*. [47] have just published a graph-neural network-based method trained on structural data as a general purpose model that can serve for further finetuning on a variety of tasks. Another concurrent work was proposed by [63] who used a geometrically variant of the perceptron trained on the largest amount of structural data to date and that leads to unprecedented results.

On the other hand, our method is based on a standard BERT architecture on which additionnal embedding modules are added to encode the structural data. This approach appeared to us simpler and more natural when we started to design our algorithm : it rely on the solid foundation provided by the research conducted on the NLP framework and our IPF solver elegantly degenerates to the classic BERT model when the additionnal embeddings modules are cut out. As such, it can directly be compared to standard NLP architecture and be easily implemented within any standard NLP-transformer based framework. Finally, our approach allows to design more proficient models using more recent transformer architecture (elektra [30], performer [32], etc.)

## III. Model

In this work, we share the same interest brought by the protein folding breakthrough and contribute by proposing a simple model that joins NLP-inspired sequenced-based model and structure encoding models to solve the inverse folding problem. This problem can be formulated as inferring the probability *p*(*t*_*n*_|**c**, *t*_1_, …*t*_*n*−1_, *t*_*n*+1_, …, *t*_*N*_) where **c** = [**c**_*n,i*_] are the 3D coordinates of every atom *i* of the *n*-th residue of the protein structure and **t** = [*t*_1_, *t*_2_, …, *t*_*N*_] is the primary sequence of the protein and where every *t*_*n*_ takes its value in Σ = {*A, C, D, E, F, G, H, I, K, L, M, N, P, Q, R, S, T, V, W, Y*} the dictionary of amino acids.

The main idea behind PeTriBERT is to consider that the inverse folding problem and the BERT-based NLP solution are strongly related: the Mask-language modeling task can be considered as a particular case of the inverse protein folding problem where the structure is unknown *i*.*e* inferring the marginal distribution *p*(*t*_*n*_|*t*_1_, …*t*_*n*−1_, *t*_*n*+1_, …, *t*_*N*_). Conversely, the classical BERT setting on the masked-language proxy task intuitively calls for structure-related feature augmentation.

To achieve this, we treat proteins as sentences and residues as words/tokens from an NLP framework point of view as in former contributions [9], [10], [12] and encode protein data using a light BERT model [16]. The main difference with pure NLP models is the use of additionnal embeddings to feed the model with tridimensional information in a way inspired by [17] and illustrated by figure 1. Indeed, besides the classic token embedding and positional embedding described in the original transformer article [64], we incorporate two additional embedding modules followed by dense layers.

**Fig. 1:**
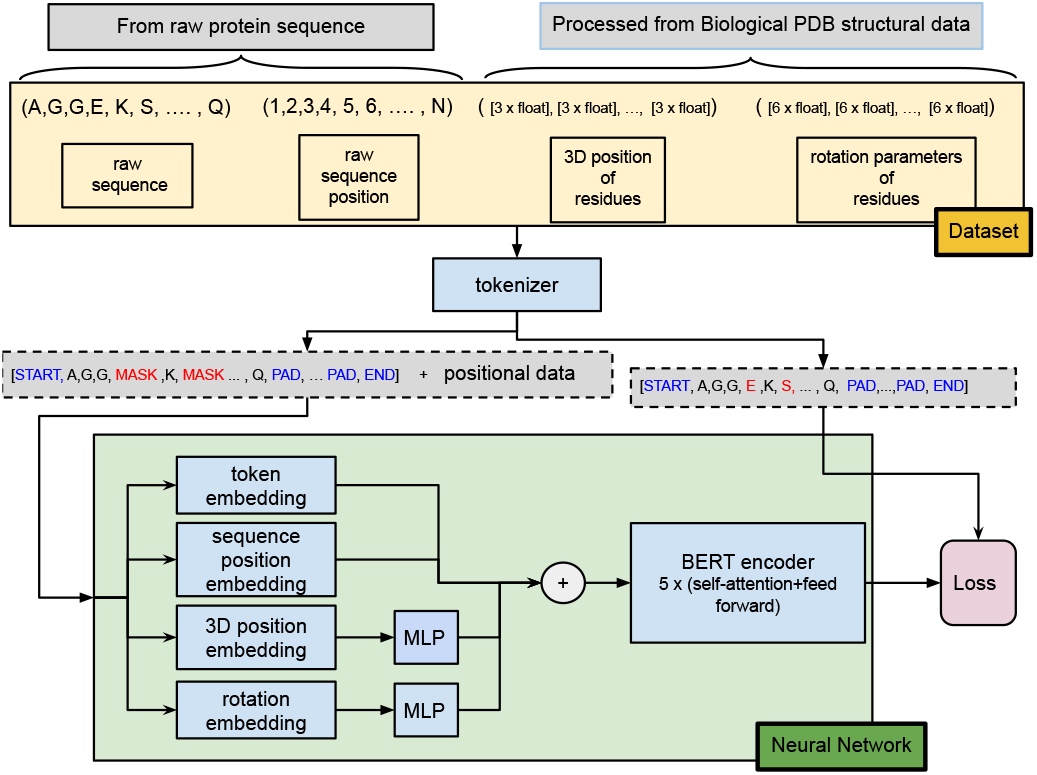
Model architecture. The architecture of PeTriBERT is based on the standard BERT with additional embedding modules: *3D position embedding* and *rotation embedding* whose role is to capture structural elements, as provided by AlphaFold for the training data. The models is trained to predict the amino-acid sequence, in a classical masked language modeling setup.

### A. Tokenization of the primary sequence

Every protein is first tokenized and then truncated or padded to a fixed length *M*. The resulting sequences include additional start, end and padding tokens, that are appended to form a new sequence **u** = ⟨*u*_1_, *u*_2_, …, *u*_*M*_ ⟩ defined by:

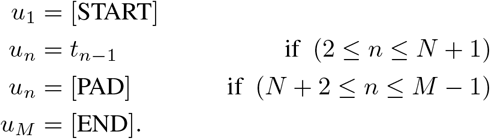

The obtained sequences **u** are used as training labels while training input sequences are obtained using random replacement or masking with the a special token [MASK] following the exact same rule as BERT [16], *i*.*e* by randomly replacing/masking 15 % of the tokens of the sequence^1^.

In the following we remove the index *n* denoting the position of the residue when unnecessary.

### B. Pre-processing of the protein structure data

Along with the protein primary sequences, a summary of the protein structure data is used as additional positional inputs by the model. In AlphafoldDB, the structural data is given as 3D coordinates for every atom comprising each residue. Before training, we preprocess this large amount of data to extract structural information for each residue from each protein in the data, composed of a single set of parameters per residue: centroid and rotation, instead of the original information at the atomic level.

1. We compute the 3D coordinates of each residue as the mean along the 3 dimensions of every of its atom:

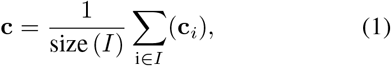

where **c** is the 3D vector of coordinates of the residue, **c**_*i*_ is the 3D vector of the *i*-th atom of the residue and *I* is the subset of atoms contained in the residue.
2. We extract rotation parameters for each residue. These parameters are obtained from the matrix **O** made of the orthogonal and normal eigenvectors of the spatial covariance matrix of each residue atomic data and verifies:

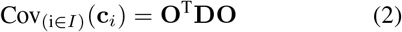

where **D** is the diagonal matrix of eigenvalues of Cov_(i∈*I*)_(**c**_*i*_)

To further compress the rotation data, we use a parametrization of the matrix **O** by solving:

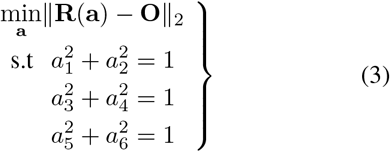

Where **a** = [*a*_1_, *a*_2_, *a*_3_, *a*_4_, *a*_5_, *a*_6_]^T^ is the vector of parameters of the rotations of the residue and **R**(**a**) is a rotation matrix parametrized over **a**. The rotation matrix **R**(**a**) is parametrized as a product of 3 rotation matrices, each being a rotation around a different axis:

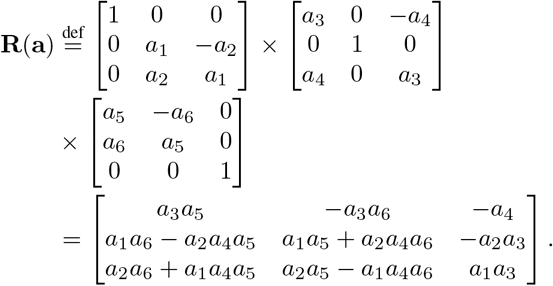

The constraints of the optimization problem force the parameters *a*_1_, *a*_2_, *a*_3_, *a*_4_, *a*_5_ and *a*_6_ to be homogeneous to cosine and sine of the angle of each of the rotations. Also note that the solution of the **O** matrix derived from the diagonalization procedure is not unique but any direct orthonormal^2^ solution is considered acceptable in our model and we think the lack of unicity of the solution is even beneficial as a form of data augmentation.

### C. Embedding Modules

The pre-processed structure data is passed to the embedding modules along with the classic uni-dimensional position and tokens. In order to make this article self-contained and help the reader comprehension, we describe in the following both the uni-dimensional positional encoding formulation as described in the original transformer [64] and the generalisation proposed for multi-dimensional positional encoding described by [17] we use to encode the structure data.

#### 1) Uni-dimensional positional encoding of the absolute position

Uni-dimensional embedding is performed by using a fixed fourier Basis functions to encode the positions in a *D*-dimensional vector. This is achieved first by computing the product

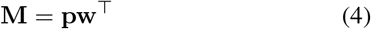

where **p** = [1, 2, …, *N*] is the *N* × 1 vector of absolute positions of the sequence and **w** is the 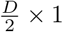 vector with

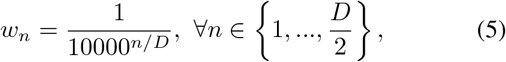

and then, by taking the cosine and sine of the matrix and concatenating:

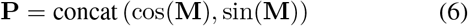

where concat designate the concatenation operation along the second dimension of the matrices. The obtained encoding *N* × *D* matrix **P** is equivalent to the original formulation in [64].

#### 2) Learnable multi-dimensional positional encoding of the 3D coordinates and rotations

To encode the 3D coordinates and the rotation parameters obtained in section III-B of every residues of the sequence we use fourier Basis functions. For the multi-dimensional case though, we encode a *L*-dimensional position matrix **P**_mult_ of size *N* × *L*, by defining a 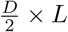 matrix **W**_*mult*_ that is randomly initialized and then learned during the training of the network. Then, we compute the product of the matrices like in the uni-dimensional case:

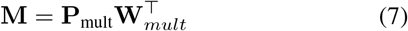

and we build a multi-dimensionnal embedding using Eq. (6). This achieves learnable random fourier features that have been shown to approximate a kernel on the euclidian distance [17]. In addition, we add a multi-layer perceptron after the encoding to improve the expressiveness of the encoding.

### D. Data Augmentation

Another contribution in this article, is the use of data augmentation to prevent over-fitting. Two types of data augmentation are performed on the structural data:

#### 1) Rigid transformation augmentation

The representation of the structural data requires to be invariant to rigid transformation of the proteins coordinates *i*.*e* be invariant to rotations and translation of the complete protein structure. For that reason and during training, we augment the coordinates on per protein basis by randomly drawing a translation vector and a rotation matrix applied globally to the coordinates of the residues.

#### 2) Non-rigid transformation augmentation

To prevent our model to capture the inherent modeling errors of the structure data estimated by AlphaFold results or to simply just memorize the training data, we add a random Gaussian noise both on the coordinates and rotation features of the inputs. The standard deviation of that random augmentation noise is typically chosen to be of the same order of magnitude of the smallest distance observed between residues for the corresponding feature for location parameters, and chosen to achieve a deviation of about 30 degree for rotation features.

## IV. Training

### A. Setting

#### 1) Hardware

Training was performed on one computing node with 2 Intel Cascade Lake 6226 processors (2 × 12 cores at 2.7 GHz) and 8 GPU Nvidia Tesla V100 SXM2 32Go during 70 hours for every variant of the model with a batch size of 128 sequences (sub-batch of 16 sequences per GPU).

#### 2) Dataset

We trained our model using the AlphaFold Dataset who was made of about 360 000 proteins at the time of our experiment. We used 80% for training (about 290 000 proteins), 10 % for validation (about 36 500 proteins) and 10 % for test (about 36 500 proteins). The AlphaFoldDB dataset was pre-processed according to section III-B. Within the model, input sequences are truncated or padded to a length of 1024 and tokenized according to section III-A.

#### 3) Model setting and Hyper-parameters

We use a small BERT configuration of 5 layers, 12 attention heads and query dimensions of 64. The feedforward part of each transformer block use a hidden size of 3072. The total amount of trainable parameter reach no more than 40 millions.

#### 4) Optimization and hyper-parameter search

The training of the model was performed using the AdamW optimizer [65] with gradient clipping and using a linear decay scheduling with linear warmup. We used the following hyper-parameters for training:

- Maximal learning rate: 1e-3
- Ending learning rate: 1e-7
- Number of warmup batches: 30 000
- Number of batches before ending rate: 250 000
- *β*_1_ =0.9, *β*_2_ =0.99,
- weight decay = 0.01.

The learning rate, number of warm-up steps and scheduler decay slope were selected using parameter sweeping. Validation loss and accuracy were logged every 100 batches.

#### 5) Data augmentation hyper-parameters

Data augmentation was performed according to section III-D. The applied rigid translation was drawn to follow an uniform distribution of 20 angström along the x, y and z axis and the applied rigid transformation was drawn to to follow an uniform distribution on the unity circle. The standard deviation of translation noise was chosen to be of 1 angström magnitude and the rotation noise to perform a random rotation of 10 degree on average.

### B. Comparing positional embedding types

In order to determine the efficiency of incorporating structural data with multi-dimensional embedding, we tried 9 different variants of our model consisting in keeping or removing embedding modules. This can be though as an ablation study as we tried to determine how each modality (position in sequence, position of residues, rotations of residues) were impacting the results. In the following, we use the following naming convention:

- **uni** in the variant name: variants where the unidimensional positional embedding of the position of the sequence is used. These variants contains the classic NLP embedding of the absolute position of residues of the sequence as described in the original transformer [64].
- **tri** in the variant name: variants where the tri-dimensional positional embedding of the coordinates of the residues (mean of the coordinates of the atoms of residues, see section III-B) is used.
- **rot** in the variant name: variants where the 6-dimensional positional embedding of the rotations of the residues (eigen vectors cosine and sin angles of the coordinates of the atoms of residues, see section III-B) is used.
- **MLP** in the variant name: an small multi-layer perceptron is added behind the residues coordinates embedding module or residues rotation embedding module if present in the variant.

We tried the 9 following variants: **(1) uni, (2) tri + rot + MLP, (3) tri + rot, (4) tri + MLP, (5) tri, (6) uni + tri + rot + MLP, (7) uni + tri + rot, (8) uni + tri + MLP, (9) uni + tri**.

Figures 2, 3 and 4 respectively show the train loss, the validation loss and the validation accuracy observed during training. Table I provides the loss and accuracy of the test dataset for the 9 variants.

**Fig. 2:**
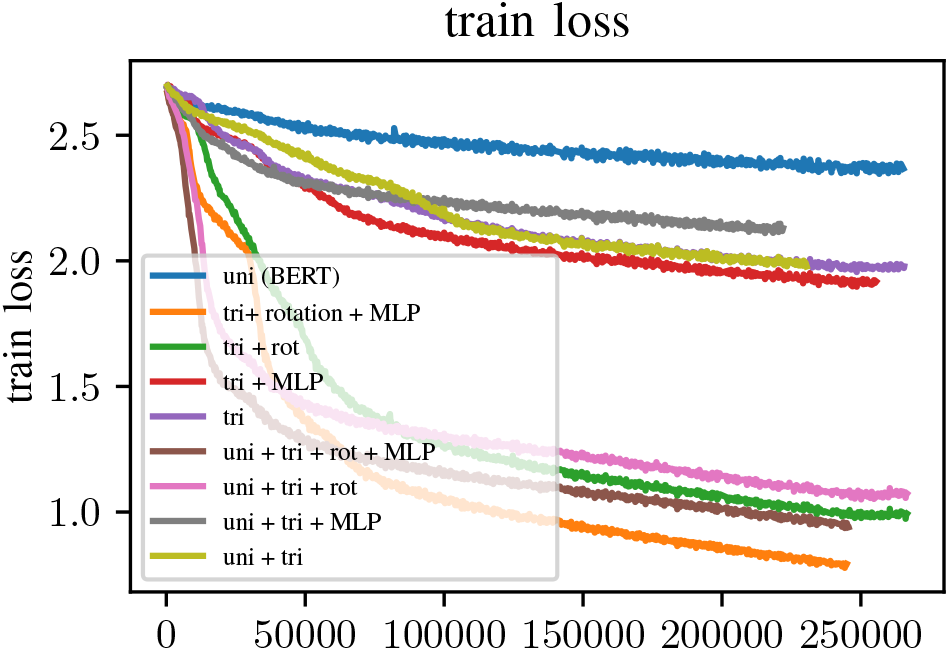
train loss across time for all 9 tested variants.

**Fig. 3:**
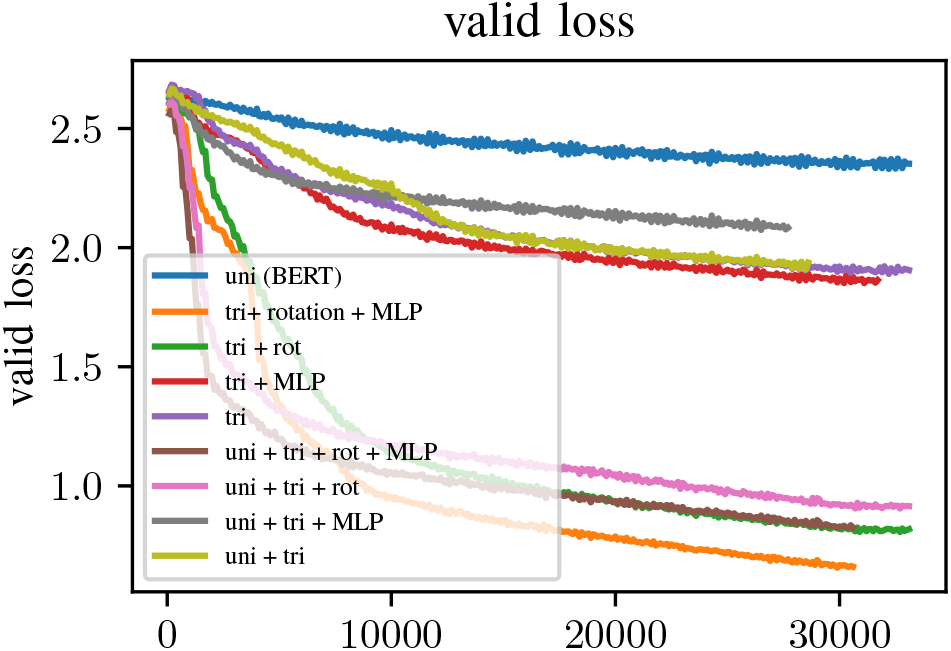
valid loss across time for all 9 tested variants.

**Fig. 4:**
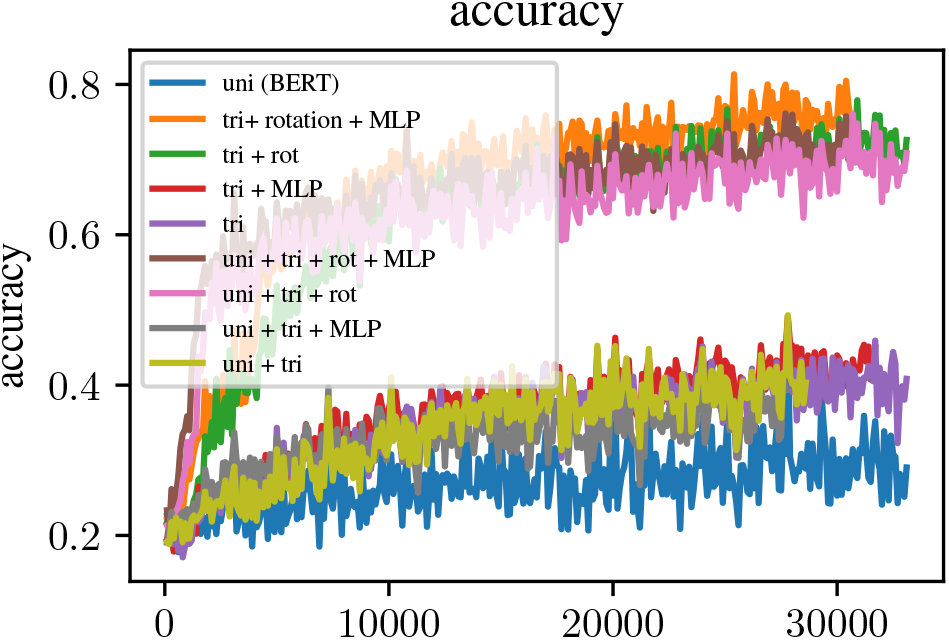
valid accuracy across time for all 9 tested variants.

**TABLE I:**
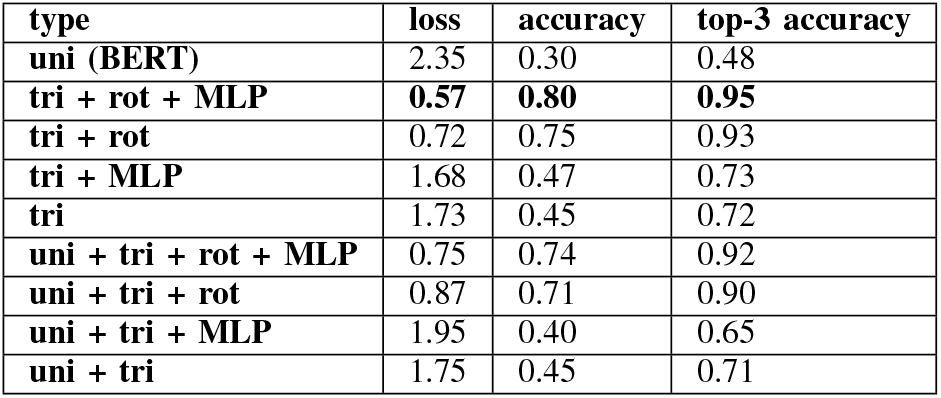
Comparison between different types of positional embedding. Loss, accuracy and top3 accuracy on the test dataset for the 9 variant of the PeTriBERT model. The overall best variant is ‘tri + rot + MLP’ where the model encodes the protein structures followed by small multilayer perceptrons.

From the results, it appears that the base BERT model performs worse than the other variants which highlights the difficulties of single-sequence model to capture the structural information from the primary sequence alone. This was verified both during training (Figures 2, 3) but also at test time I. Amongst the 8 remaining variants, it appears that residue rotation fed methods have a clear edge over the methods that don’t use this type of embedding. At test time, accuracy for these methods is on average about 0.25 points higher than the rest of the methods. More surprisingly, methods that include the classic BERT unidirectional encoder tends to perform worse than their non-unidirectional counterpart: for instance the ‘uni + tri + rot’ variant showed an accuracy of 0.71 against 0.75 for the ‘tri + rot variant’. Although one may think there is arguably no good reason for a deep learning model to perform worse when fed with more information, one possible explanatory reason relies in the huge informative gap between the primary sequence data and the structural data: considering our model is very small compared to modern architectures featuring only 40M parameters, we believe that the uni-dimensional embedding are regarded as noise by the model and therefore worsen the ability of our model to accurately predict missing residues. Finally, the MLP enhanced embedding variants display some improvement over the non-MLP enhanced one, highlighting the complexity of encoding the protein structure and capture the geometrical patterns.

## V. PROTEIN GENERATION

### A. Generative method

PeTriBERT is aimed at solving the inverse folding problem and as such can be used for protein generation. We achieve this by using Gibbs sampling, inspired by [18], [19]. The goal is to generate a novel protein sharing the same protein structure than a given target. To perform generation, we initialize the algorithm by retrieving the target protein, its primary sequence and structure data (known or estimated with Alphafold). Then, we perform a loop over each residue (denoted *n* in the following) of the sequence where:

1. we mask the current amino acid *n* data in the primary sequence,
2. we feed the primary sequence and structure data to PeTriBERT that returns a predictive distribution *p*(*t*_*n*_|*t*_1_, …*t*_*n*−1_, *t*_*n*+1_, …*t*_*N*_, *c*) given the other amino acids *p*(*t*_*n*_|*t*_1_, …*t*_*n*−1_, *t*_*n*+1_, …*t*_*N*_) and structure data *c*.
3. We sample from this distribution and replace *t*_*i*_ with the results in the sequence

This loop is repeated many times, each one of them being initialized with the output of the previous one. Also, at the end of every loop, we store the obtained candidate sequence.

When the generation is over, we send all the generated sequences to AlphaFold and select the one achieving the top lDDT score [66].

### B. In silico generation of avGFP protein variants

To assess our method, we tested it using the green fluorescent protein from Aequorea victoria (avGFP) [20] as target structure, as it is likely one of the best studied protein so far with a very recognizable barrel structure. We performed 2 different experiments using Gibbs sampling:

- Generation of 200 hundred proteins using avGFP sequences as target structure exactly as described in section V-A.
- Generation of 200 hundred proteins using avGFP sequences where the amino acid of position 65, 66 and 67 are forced to have the original sequence residues, respectively S (Serine), Y (Tyrosine) and G (Glycine). This was enforced as these residues are known to be essential to the fluorescence properties of the protein.

When the algorithm terminates is over, the 400 generated protein sequences are processed through AlphaFold, and we compute the lDDT score between each candidate against the avGFP protein of reference.

Despite using a very simple Gibbs sampling scheme, the algorithm often outputs sequence featuring qualitatively similar protein structure apart for a few iteration where the folding process seems incomplete as only chunks of the structure folds accordingly to the query as depicted on figure. 5.

**Fig. 5:**
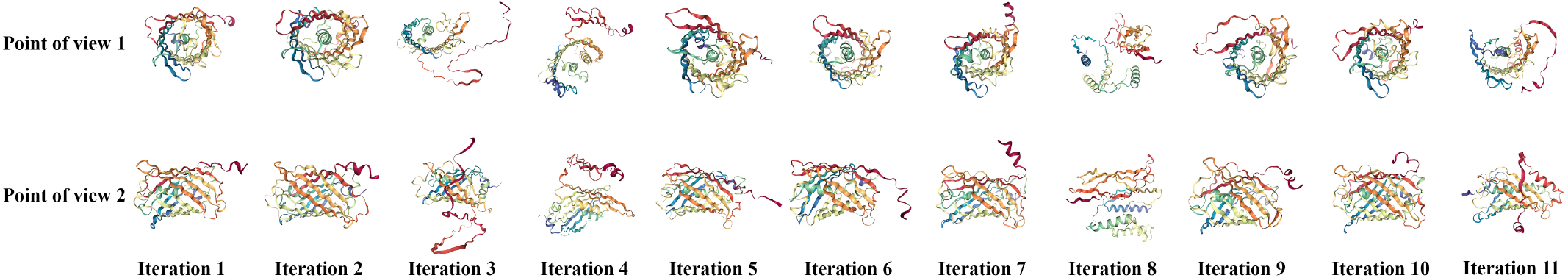
Computation history of the generated proteins. This figure presents the structure predicted by alphafold at every iteration of our gibbs sampling generator powered by PeTriBERT.

However, if we keep the five best scoring proteins of each group, we obtain proteins displaying a score ranging from 0.70 to 0.79 and a similarity in sequence of about 20% for each as shown in Table V. When blasted on the whole proteome 9 of 10 synthetic GFP returns no homologues. This shows that PetriBert is able to design purely synthetic proteins with a given 3D structure as validated by IDDT score. Biosynthesis of these synthetic GFP variants is a work in progress. Figure 6 gives a comparison between the best scoring protein and the target.

**TABLE II:** Precision and recall per residue of tri + MLP.

| residue | precision | recall | residue | precision | recall |
| --- | --- | --- | --- | --- | --- |
| **A** | 0.49 | 0.49 | **M** | 0.60 | 0.24 |
| **C** | 0.66 | 0.40 | **N** | 0.43 | 0.35 |
| **D** | 0.46 | 0.46 | **P** | 0.52 | 0.75 |
| **E** | 0.41 | 0.53 | **Q** | 0.44 | 0.25 |
| **F** | 0.55 | 0.36 | **R** | 0.70 | 0.55 |
| **G** | 0.67 | 0.67 | **S** | 0.36 | 0.52 |
| **H** | 0.53 | 0.24 | **T** | 0.42 | 0.28 |
| **I** | 0.44 | 0.29 | **V** | 0.40 | 0.38 |
| **K** | 0.43 | 0.53 | **W** | 0.73 | 0.34 |
| **L** | 0.41 | 0.61 | **Y** | 0.63 | 0.39 |

**TABLE III:** Precision and recall per residue of tri + rot + MLP.

| residue | precision | recall | residue | precision | recall |
| --- | --- | --- | --- | --- | --- |
| **A** | 0.95 | 0.96 | **M** | 0.74 | 0.60 |
| **C** | 0.80 | 0.70 | **N** | 0.69 | 0.65 |
| **D** | 0.74 | 0.73 | **P** | 0.97 | 0.98 |
| **E** | 0.71 | 0.78 | **Q** | 0.64 | 0.50 |
| **F** | 0.73 | 0.76 | **R** | 0.81 | 0.72 |
| **G** | 0.99 | 0.99 | **S** | 0.85 | 0.88 |
| **H** | 0.69 | 0.65 | **T** | 0.75 | 0.72 |
| **I** | 0.79 | 0.79 | **V** | 0.75 | 0.80 |
| **K** | 0.71 | 0.81 | **W** | 0.90 | 0.80 |
| **L** | 0.82 | 0.87 | **Y** | 0.71 | 0.65 |

**TABLE IV:**
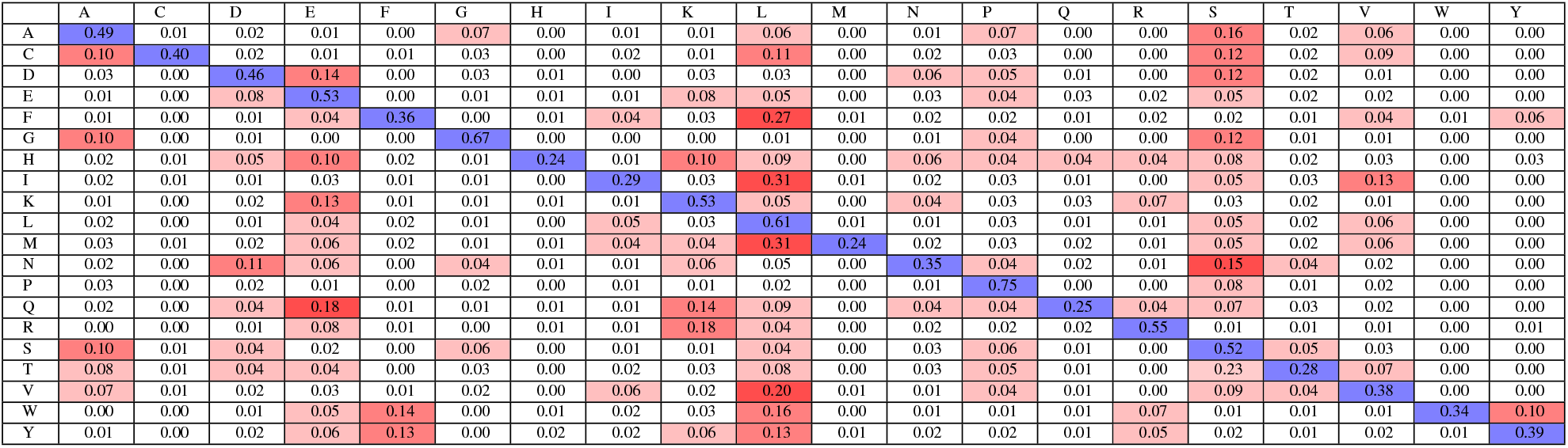
Confusion matrix of tri + MLP variant for each residue. Rows represent groundtruth and columns represent predictions. Values are given normalized over rows so that every cell represent the prediction rate knowing the ground-truth and therefore blue cells gives recall. For instance, when a residue of type C should be given, 10 % of the time a residue of type A is predicted.

**TABLE V:** lDDT Score and rate of similary of generated proteins against avGFP. gen0, gen1, gen4, gen55 and gen90 are respectively the protein generated at iteration 0, 1, 4, 55 and 90 of the first generation group. genSYG69, genSYG117, genSYG126, genSYG141 and genSYG174 are respectively the protein generated at iteration 69, 117, 126, 141 and 174 of the second group, where the 3 residues at position 65, 66, and 67 are enforced to be S, Y and G.

| generated sequence | IDDT score | homology rate |
| --- | --- | --- |
| gen0 | 0.74 | 50 (21%) |
| gen1 | 0.70 | 44 (18%) |
| gen4 | 0.77 | 43 (18%) |
| gen55 | 0.72 | 43 (18%) |
| gen90 | 0.75 | 46 (19%) |
| genSYG69 | 0.76 | 42 (17%) |
| genSYG117 | 0.79 | 45 (18%) |
| genSYG126 | 0.75 | 44 (18%) |
| genSYG141 | 0.72 | 49 (20%) |
| genSYG174 | 0.76 | 44 (18%) |

**Fig. 6:**
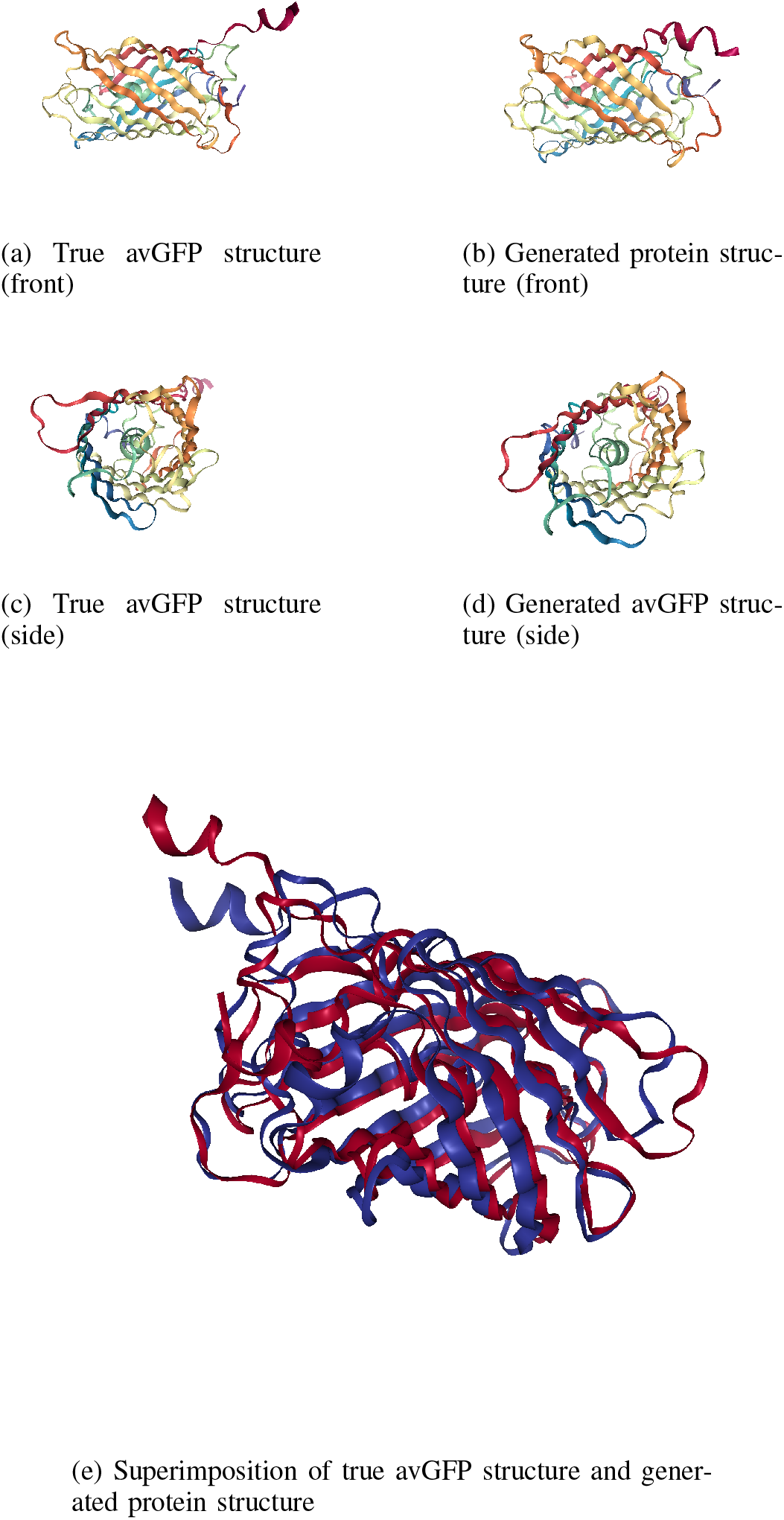
3D plots of the avGFP structure and the generated structure predicted by AlphaFold. (6a) and (6c) display the true avGFP structure seen from two different angles, while (6b) and (6d) display one of our top scoring generated protein from the same angles. (6e) display the superimposition of both reference and generated structure.

## VI. Discussion

In this article, we proposed PeTriBERT, a simple transformer architecture that aims to solve the inverse folding protein problem and used it as basis to perform *de novo* protein generation from a structure constraint. To achieve this, PeTriBERT replaces the usual positional encoding of the BERT model by a multi-dimensional spatial encoding of the protein structure and benefits from the recent advances in protein folding. We found out that PeTriBERT is able to generate good quality novel protein despite being considerably smaller than modern neural networks, using only 40 M parameters, highlighting the capacity of out-of-the-box architecture to distillate the byproducts of protein folding.

Recently, a similar work taking benefits from the structural output of AlphaFold 2 has been published [63]. Compared to our work, this model is auto-regressive (our is an auto-encoder), is based on a complex adhoc architecture [6], uses more parameters (142 M for the largest version against 40 for our) and learned on much more Data (12M proteins agains around 350 000 for our).

At the opposite, our model is more simple, relates to the well-know NLP framework and easy to implement as it relies on a simple trick that consists in i) using a tridimensional positionnal encoding of a transformer encoder ii) data augmentation to train the network to be rotation and translation invariant. As such, we hope our model can give some insights on a pratical way to incorporate the structural data of recent forlding technics to transformers-based network.

Nevertheless, it appears that such models could highly benefit from using more parameters and larger datasets, which will be feasible as the part of the proteome that provide structure data will very likely grow over the few next years. Together with ongoing experimental validations, we are now implementing such PetriBert extentions for following transfer learning studies.

## Acknowledgment

This work was granted access to the HPC resources of IDRIS under the allocation 20XX-101719 made by GENCI. This research is supported by MUSE University of Montpellier research grant (AI3P: Artificial Intelligence to Predict Phenotypes in Plants), the ANR (the French National Research Agency) under the “Investissements d’avenir” programme with the reference ANR-16-IDEX-0006 to GK.

## Footnotes

1 Or more exactly, we randomly select 15% of the tokens of the sequences. Among the selection, 80% are replaced with the [MASK] token, 10% are randomly replaced, and 10% are left unchanged.

2 After the diagonalization procedure, if the obtained set of vectors form an non-direct orthonormal basis, we invert one of the obtained vector.

## Notes

### Competing Interest Statement

The authors have declared no competing interest.

## Bibliography

[1] C. Pabo, “Molecular technology: designing proteins and peptides,” Nature, vol. 301, no. 5897, pp. 200–200, 1983.

[2] A. Leaver-Fay, M. Tyka, S. M. Lewis, O. F. Lange, J. Thompson, R. Jacak, K. W. Kaufman, P. D. Renfrew, C. A. Smith, W. Sheffler et al., “Rosetta3: an object-oriented software suite for the simulation and design of macromolecules,” in Methods in enzymology. Elsevier, 2011, vol. 487, pp. 545–574.

[3] S. Min, B. Lee, and S. Yoon, “Deep learning in bioinformatics,” Briefings in bioinformatics, vol. 18, no. 5, pp. 851–869, 2017.

[4] A. Strokach, D. Becerra, C. Corbi-Verge, A. Perez-Riba, and P. M. Kim, “Fast and flexible protein design using deep graph neural networks,” Cell systems, vol. 11, no. 4, pp. 402–411, 2020.

[5] J. Ingraham, V. Garg, R. Barzilay, and T. Jaakkola, “Generative models for graph-based protein design,” Advances in Neural Information Processing Systems, vol. 32, 2019.

[6] B. Jing, S. Eismann, P. Suriana, R. J. Townshend, and R. Dror, “Learning from protein structure with geometric vector perceptrons,” arXiv preprint 2009.01411, 2020.

[7] V. Gligorijević, P. D. Renfrew, T. Kosciolek, J. K. Leman, D. Berenberg, T. Vatanen, C. Chandler, B. C. Taylor, I. M. Fisk, H. Vlamakis et al., “Structure-based protein function prediction using graph convolutional networks,” Nature communications, vol. 12, no. 1, pp. 1–14, 2021.

[8] F. Baldassarre, D. Menéndez Hurtado, A. Elofsson, and H. Azizpour, “Graphqa: protein model quality assessment using graph convolutional networks,” Bioinformatics, vol. 37, no. 3, pp. 360–366, 2021.

[9] A. Rives, J. Meier, T. Sercu, S. Goyal, Z. Lin, J. Liu, D. Guo, M. Ott, C. L. Zitnick, J. Ma, and R. Fergus, “Biological structure and function emerge from scaling unsupervised learning to 250 million protein sequences,” Proceedings of the National Academy of Sciences, vol. 118, no. 15, Apr. 2021, publisher: National Academy of Sciences Section: Biological Sciences. [Online]. Available: https://www.pnas.org/content/118/15/e2016239118

[10] R. Rao, J. Meier, T. Sercu, S. Ovchinnikov, and A. Rives, “Transformer protein language models are unsupervised structure learners,” p. 24.

[11] A. Elnaggar, M. Heinzinger, C. Dallago, G. Rihawi, Y. Wang, L. Jones, T. Gibbs, T. Feher, C. Angerer, M. Steinegger, D. Bhowmik, and B. Rost, “ProtTrans: Towards Cracking the Language of Life ‘s Code Through Self-Supervised Deep Learning and High Performance Computing,” 2007.06225 [cs, stat], May 2021, 2007.06225. [Online]. Available: http://arxiv.org/abs/2007.06225

[12] A. Nambiar, S. Liu, M. Hopkins, M. Heflin, S. Maslov, and A. Ritz, “Transforming the Language of Life: Transformer Neural Networks for Protein Prediction Tasks,” bioRxiv, p. 2020.06.15.153643, Jun. 2020, publisher: Cold Spring Harbor Laboratory Section: New Results. [Online]. Available: https://www.biorxiv.org/content/10.1101/2020.06.15.153643v1

[13] J. Jumper, R. Evans, A. Pritzel, T. Green, M. Figurnov, O. Ronneberger, K. Tunyasuvunakool, R. Bates, A. Žídek, A. Potapenko, A. Bridgland, C. Meyer, S. A. A. Kohl, A. J. Ballard, A. Cowie, B. Romera-Paredes, S. Nikolov, R. Jain, J. Adler, T. Back, S. Petersen, D. Reiman, E. Clancy, M. Zielinski, M. Steinegger, M. Pacholska, T. Berghammer, S. Bodenstein, D. Silver, O. Vinyals, A. W. Senior, K. Kavukcuoglu, P. Kohli, and D. Hassabis, “Highly accurate protein structure prediction with AlphaFold,” Nature, vol. 596, no. 7873, pp. 583–589, Aug. 2021. [Online]. Available: https://www.nature.com/articles/s41586-021-03819-2

[14] M. Baek, F. DiMaio, I. Anishchenko, J. Dauparas, S. Ovchinnikov, G. R. Lee, J. Wang, Q. Cong, L. N. Kinch, R. D. Schaeffer et al., “Accurate prediction of protein structures and interactions using a three-track neural network,” Science, vol. 373, no. 6557, pp. 871–876, 2021.

[15] M. Varadi, S. Anyango, M. Deshpande, S. Nair, C. Natassia, G. Yordanova, D. Yuan, O. Stroe, G. Wood, A. Laydon, A. Žídek, T. Green, K. Tunyasuvunakool, S. Petersen, J. Jumper, E. Clancy, R. Green, A. Vora, M. Lutfi, M. Figurnov, A. Cowie, N. Hobbs, P. Kohli, G. Kleywegt, E. Birney, D. Hassabis, and S. Velankar, “AlphaFold Protein Structure Database: massively expanding the structural coverage of protein-sequence space with high-accuracy models,” Nucleic Acids Research, no. gkab1061, Nov. 2021. [Online]. Available: https://doi.org/10.1093/nar/gkab1061

[16] J. Devlin, M.-W. Chang, K. Lee, and K. Toutanova, “Bert: Pre-training of deep bidirectional transformers for language understanding,” arXiv preprint 1810.04805, 2018.

[17] Y. Li, S. Si, G. Li, C.-J. Hsieh, and S. Bengio, “Learnable fourier features for multi-dimensional spatial positional encoding,” Advances in Neural Information Processing Systems, vol. 34, 2021.

[18] A. Wang and K. Cho, “Bert has a mouth, and it must speak: Bert as a markov random field language model,” arXiv preprint 1902.04094, 2019.

[19] S. R. Johnson, S. Monaco, K. Massie, and Z. Syed, “Generating novel protein sequences using gibbs sampling of masked language models,” bioRxiv, 2021.

[20] D. C. Prasher, V. K. Eckenrode, W. W. Ward, F. G. Prendergast, and M. J. Cormier, “Primary structure of the aequorea victoria green-fluorescent protein,” Gene, vol. 111, no. 2, pp. 229–233, 1992.

[21] Y. Cao, P. Das, V. Chenthamarakshan, P.-Y. Chen, I. Melnyk, and Y. Shen, “Fold2seq: A joint sequence (1d)-fold (3d) embedding-based generative model for protein design,” in International Conference on Machine Learning. PMLR, 2021, pp. 1261–1271.

[22] G. Derevyanko, S. Grudinin, Y. Bengio, and G. Lamoureux, “Deep convolutional networks for quality assessment of protein folds,” Bioinformatics, vol. 34, no. 23, pp. 4046–4053, 2018.

[23] B. Rozemberczki, S. Bonner, A. Nikolov, M. Ughetto, S. Nilsson, and E. Papa, “A Unified View of Relational Deep Learning for Drug Pair Scoring,” 2111.02916 [cs], Nov. 2021, 2111.02916. [Online]. Available: http://arxiv.org/abs/2111.02916

[24] T. Mikolov, K. Chen, G. Corrado, and J. Dean, “Efficient estimation of word representations in vector space,” arXiv preprint 1301.3781, 2013.

[25] E. Asgari and M. R. Mofrad, “Continuous distributed representation of biological sequences for deep proteomics and genomics,” PloS one, vol. 10, no. 11, p. e0141287, 2015.

[26] D. Kimothi, A. Soni, P. Biyani, and J. M. Hogan, “Distributed representations for biological sequence analysis,” arXiv preprint 1608.05949, 2016.

[27] C. Mazzaferro, “Predicting protein binding affinity with word embeddings and recurrent neural networks,” bioRxiv, p. 128223, 2017.

[28] P. Ng, “dna2vec: Consistent vector representations of variable-length k-mers,” arXiv preprint 1701.06279, 2017.

[29] Y. Liu, M. Ott, N. Goyal, J. Du, M. Joshi, D. Chen, O. Levy, M. Lewis, L. Zettlemoyer, and V. Stoyanov, “Roberta: A robustly optimized bert pretraining approach,” arXiv preprint 1907.11692, 2019.

[30] K. Clark, M.-T. Luong, Q. V. Le, and C. D. Manning, “Electra: Pretraining text encoders as discriminators rather than generators,” arXiv preprint 2003.10555, 2020.

[31] N. Kitaev, Ł. Kaiser, and A. Levskaya, “Reformer: The efficient transformer,” arXiv preprint 2001.04451, 2020.

[32] K. Choromanski, V. Likhosherstov, D. Dohan, X. Song, A. Gane, T. Sarlos, P. Hawkins, J. Davis, A. Mohiuddin, L. Kaiser et al., “Rethinking attention with performers,” arXiv preprint 2009.14794, 2020.

[33] A. Jaegle, F. Gimeno, A. Brock, O. Vinyals, A. Zisserman, and J. Carreira, “Perceiver: General perception with iterative attention,” in International Conference on Machine Learning. PMLR, 2021, pp. 4651–4664.

[34] M. Heinzinger, A. Elnaggar, Y. Wang, C. Dallago, D. Nechaev, F. Matthes, and B. Rost, “Modeling the language of life–deep learning protein sequences,” Biorxiv, p. 614313, 2019.

[35] M. E. Peters, M. Neumann, M. Iyyer, M. Gardner, C. Clark, K. Lee, and L. Zettlemoyer, “Deep contextualized word representations,” CoRR, vol. abs/1802.05365, 2018. [Online]. Available: http://arxiv.org/abs/1802.05365

[36] N. Brandes, D. Ofer, Y. Peleg, N. Rappoport, and M. Linial, “ProteinBERT: A universal deep-learning model of protein sequence and function,” Bioinformatics, preprint, May 2021. [Online]. Available: http://biorxiv.org/lookup/doi/10.1101/2021.05.24.445464

[37] J. Vig, A. Madani, L. R. Varshney, C. Xiong, R. Socher, and N. F. Rajani, “Bertology meets biology: Interpreting attention in protein language models,” arXiv preprint 2006.15222, 2020.

[38] R. Rao, N. Bhattacharya, N. Thomas, Y. Duan, P. Chen, J. Canny, P. Abbeel, and Y. Song, “Evaluating protein transfer learning with tape,” Advances in neural information processing systems, vol. 32, 2019.

[39] R. M. Rao, J. Liu, R. Verkuil, J. Meier, J. Canny, P. Abbeel, T. Sercu, and A. Rives, “Msa transformer,” in International Conference on Machine Learning. PMLR, 2021, pp. 8844–8856.

[40] C. Yanofsky, V. Horn, and D. Thorpe, “Protein structure relationships revealed by mutational analysis,” Science, vol. 146, no. 3651, pp. 1593–1594, 1964.

[41] D. Altschuh, A. Lesk, A. Bloomer, and A. Klug, “Correlation of co-ordinated amino acid substitutions with function in viruses related to tobacco mosaic virus,” Journal of molecular biology, vol. 193, no. 4, pp. 693–707, 1987.

[42] D. Altschuh, T. Vernet, P. Berti, D. Moras, and K. Nagai, “Coordinated amino acid changes in homologous protein families,” Protein Engineering, Design and Selection, vol. 2, no. 3, pp. 193–199, 1988.

[43] S. Biswas, G. Khimulya, E. C. Alley, K. M. Esvelt, and G. M. Church, “Low-n protein engineering with data-efficient deep learning,” Nature methods, vol. 18, no. 4, pp. 389–396, 2021.

[44] J. Meier, R. Rao, R. Verkuil, J. Liu, T. Sercu, and A. Rives, “Language models enable zero-shot prediction of the effects of mutations on protein function,” Advances in Neural Information Processing Systems, vol. 34, 2021.

[45] P. Hermosilla, M. Schäfer, M. Lang, G. Fackelmann, P. P. Vázquez, B. Kozlíková, M. Krone, T. Ritschel, and T. Ropinski, “Intrinsic-extrinsic convolution and pooling for learning on 3d protein structures,” arXiv preprint 2007.06252, 2020.

[46] Y. Zhang, Y. Chen, C. Wang, C.-C. Lo, X. Liu, W. Wu, and J. Zhang, “Prodconn: Protein design using a convolutional neural network,” Proteins: Structure, Function, and Bioinformatics, vol. 88, no. 7, pp. 819–829, 2020.

[47] Z. Zhang, M. Xu, A. Jamasb, V. Chenthamarakshan, A. Lozano, P. Das, and J. Tang, “Protein representation learning by geometric structure pretraining,” arXiv preprint 2203.06125, 2022.

[48] M. Ragoza, J. Hochuli, E. Idrobo, J. Sunseri, and D. R. Koes, “Protein– ligand scoring with convolutional neural networks,” Journal of chemical information and modeling, vol. 57, no. 4, pp. 942–957, 2017.

[49] H. Zeng, M. D. Edwards, G. Liu, and D. K. Gifford, “Convolutional neural network architectures for predicting dna–protein binding,” Bioinformatics, vol. 32, no. 12, pp. i121–i127, 2016.

[50] J. Jiménez, S. Doerr, G. Martínez-Rosell, A. S. Rose, and G. De Fabritiis, “Deepsite: protein-binding site predictor using 3d-convolutional neural networks,” Bioinformatics, vol. 33, no. 19, pp. 3036–3042, 2017.

[51] H. Zhang, R. Guan, F. Zhou, Y. Liang, Z.-H. Zhan, L. Huang, and X. Feng, “Deep residual convolutional neural network for protein-protein interaction extraction,” IEEE Access, vol. 7, pp. 89 354–89 365, 2019.

[52] V. R. Somnath, C. Bunne, and A. Krause, “Multi-scale representation learning on proteins,” Advances in Neural Information Processing Systems, vol. 34, 2021.

[53] K. Yue and K. A. Dill, “Inverse protein folding problem: designing polymer sequences.” Proceedings of the National Academy of Sciences, vol. 89, no. 9, pp. 4163–4167, 1992.

[54] W. Boomsma and J. Frellsen, “Spherical convolutions and their application in molecular modelling,” Advances in neural information processing systems, vol. 30, 2017.

[55] N. Anand, R. Eguchi, I. I. Mathews, C. P. Perez, A. Derry, R. B. Altman, and P.-S. Huang, “Protein sequence design with a learned potential,” Nature communications, vol. 13, no. 1, pp. 1–11, 2022.

[56] S. Chen, Z. Sun, L. Lin, Z. Liu, X. Liu, Y. Chong, Y. Lu, H. Zhao, and Y. Yang, “To improve protein sequence profile prediction through image captioning on pairwise residue distance map,” Journal of chemical information and modeling, vol. 60, no. 1, pp. 391–399, 2019.

[57] K. He, X. Zhang, S. Ren, and J. Sun, “Deep residual learning for image recognition,” in Proceedings of the IEEE conference on computer vision and pattern recognition, 2016, pp. 770–778.

[58] I. Anishchenko, S. J. Pellock, T. M. Chidyausiku, T. A. Ramelot, S. Ovchinnikov, J. Hao, K. Bafna, C. Norn, A. Kang, A. K. Bera et al., “De novo protein design by deep network hallucination,” Nature, vol. 600, no. 7889, pp. 547–552, 2021.

[59] J. Wang, S. Lisanza, D. Juergens, D. Tischer, I. Anishchenko, M. Baek, J. L. Watson, J. H. Chun, L. F. Milles, J. Dauparas et al., “Deep learning methods for designing proteins scaffolding functional sites,” bioRxiv, 2021.

[60] N. Anand and P. Huang, “Generative modeling for protein structures,” Advances in neural information processing systems, vol. 31, 2018.

[61] R. R. Eguchi, N. Anand, C. A. Choe, and P.-S. Huang, “Ig-vae: generative modeling of immunoglobulin proteins by direct 3d coordinate generation,” Biorxiv, 2020.

[62] F. Collins, The language of life: DNA and the revolution in personalised medicine. Profile Books, 2010.

[63] C. Hsu, R. Verkuil, J. Liu, Z. Lin, B. Hie, T. Sercu, A. Lerer, and A. Rives, “Learning inverse folding from millions of predicted structures,” bioRxiv, 2022.

[64] A. Vaswani, N. Shazeer, N. Parmar, J. Uszkoreit, L. Jones, A. N. Gomez, Ł. Kaiser, and I. Polosukhin, “Attention is all you need,” Advances in neural information processing systems, vol. 30, 2017.

[65] I. Loshchilov and F. Hutter, “Decoupled weight decay regularization,” arXiv preprint 1711.05101, 2017.

[66] V. Mariani, M. Biasini, A. Barbato, and T. Schwede, “lddt: a local superposition-free score for comparing protein structures and models using distance difference tests,” Bioinformatics, vol. 29, no. 21, pp. 2722–2728, 2013.

